# SLM-Transform: A Method for Memory-Efficient Indexing of Spectra for Database Search in LC-MS/MS Proteomics

**DOI:** 10.1101/531681

**Authors:** Muhammad Haseeb, Muaaz G. Awan, Alexander S. Cadigan, Fahad Saeed

**Affiliations:** School of Computing and Information Sciences, Florida International University; Department of Computer Science, Western Michigan University; Department of Computer Science, Kalamazoo College

## Abstract

The most commonly used strategy for peptide identification in shotgun LC-MS/MS proteomics involves searching of MS/MS data against an in-silico digested protein sequence database. Typically, the digested peptide sequences are indexed into the memory to allow faster search times. However, subjecting a database to post-translational modifications (PTMs) during digestion results in an exponential increase in the number of peptides and therefore memory consumption. This limits the usage of existing fragment-ion based open-search algorithms for databases with several PTMs. In this paper, we propose a novel fragment-ion indexing technique which is analogous to suffix array transformation and allows constant time querying of indexed ions. We extend our transformation method, called SLM-Transform, by constructing ion buckets that allow querying of *all* indexed ions by mass by only storing information on distribution of ion-frequencies within buckets. The stored information is used with a regression technique to locate the position of ions in constant time. Moreover, the number of theoretical b- and y-ions generated and indexed for each theoretical spectrum are limited. Our results show that SLM-Transform allows indexing of up to 4x peptides than other leading fragment-ion based database search tools within the same memory constraints. We show that SLM-Transform based index allows indexing of over 83 million peptides within 26GB RAM as compared to 80GB required by MSFragger. Finally, we show the constant ion retrieval time for SLM-Transform based index allowing ultrafast peptide search speeds.

Source code will be made available at: https://github.com/pcdslab/slmindex

## 1 Introduction

In a shotgun proteomics experiment, a complex protein mixture is proteolyzed using an enzyme, most commonly trypsin. The digested peptides are then fed to a liquid-chromatography (LC) followed by mass spectrometry (MS/MS) pipeline. The acquired tandem MS/MS spectra are identified and assigned to a peptide sequence using computational techniques. The most commonly used computational method involves searching MS/MS spectra against a protein sequence database [11]. However, a large fraction of acquired MS/MS data remain unassigned [5, 24] even using high quality data acquired from modern mass-spectrometers. Moreover, the accuracy of peptide deductions greatly depends on the parameters used for the search [20] as well as the post-processing technique used to assign confidence scores to the peptide-spectra matches (PSMs) [13].

Conventional protein database search algorithms filter peptide sequences from database based on precursor masses [9]. Hence, the spectra having unexpected mass-shifts are filtered-out and escape the search space [5, 12]. The mass-shifts in these spectra arise due to post-translational modifications (PTMs), non-specific digestion, amino acid mutations and other biological processes, collectively referred to as ‘dark matter of shotgun proteomics’ [29]. Recently, it has been shown [5, 15] that the peptide identification rate increases if speactra are searched against wide precursor mass tolerance windows of upto ±500Da (also known as open-search). However, widening the search window to such large tolerances incurs an enormous search time penalty [15, 4].

A number of computational techniques have been developed to quickly reduce the search space by filtering-in the peptide sequences that have high likelihood of being matched to a MS/MS spectrum and hence, accelerate the database search. A popular PSM filtering technique is based on sequence-tagging and is employed in [4, 18, 31, 7, 28] where a sequence-tag is extracted from the experimental spectra and its k-mers are constructed. The proteome database is filtered prior to digestion using the k-mer tags and the search is conducted only against the protein sequences containing one or more k-mer tags. However, the number of peptides generated from even a filtered proteome sequence database may still be large due to PTM’s. Other than sequence-tagging, peptide indexing has also been employed in pFind-Alioth [3], Crux [21] and Andromeda [6] to improve database search times.

Another popular indexing technique is based on database indexing and has been employed in MSFragger [15] and pFind 2.1 [16]. The *fragment-ion* indexing based techniques transform the peptide-to-spectrum cross-correlation problem into shared peak count problem. For each query spectrum, a peptide in the index is considered as a candidate peptide match if it shares more than a certain threshold number of common peaks with the query spectrum. The candidate peptides are then formally scored using a scoring algorithm [8, 25, 10] to obtain PSMs. This scoring method was boosted using parallel computing by MSFragger which sped up the search time by several hundred folds over other open-search algorithms. The purpose of open-search is to identify the MS/MS spectra containing one or more unknown modifications. For such MS/MS spectra, the correct peptide sequence is not usually present in the index, however, similar or unmodified versions of their peptide sequences can still be identified based on shared peak count. Therefore, incorporating more known PTMs implies a higher probability of presence of more similar peptides in the index which would therefore increase the probability of obtaining a higher number of correct PSMs in open-search.

However, there is a gigantic footprint (8 bytes per ion) associated with MSFragger’s Fragment-Index. MSFragger handles this by splitting its index into several chunks, running the search on each chunk separately, and combining the results. In doing so, each chunk is loaded into and discarded from the RAM during the search. If the search has to be conducted with a number of PTMs and charge states incorporated, it must be done in multiple runs to avoid a crash [23], reported at: https://github.com/chhh/FragPipe/issues/63. A similar issue was also reported by [4] where MSFragger crashed when non-tryptic peptides were included in the search space. This demands for the need of novel methods that can index large number of peptides and corresponding theoretical spectra in a memory-efficient manner as well fast peptide identification.

In this paper, we present a novel indexing technique called SLM-Transform (pronounced: *Slim Transform*) and the index constructed as a result of SLM-Transform is called SLM-Index (pronounced: *Slim Index*). The SLM-Transform is analogous to Suffix Array transformation [17] which allow constant time query for text (char) data. However, the memory footprint for Suffix Arrays is 4*x* size of input text. This limitation was later solved by construction of Burrows-Wheeler transform [2]. However, in case of fragment-ion indexing, where the data type is floating points or integers, the transformation is applied using Key-Value sort and the resultant arrays have only a minimal memory overhead over the input size (≈ 4.4 bytes per ion).

Figure 1 (A) shows application of Suffix Array transformation on a string input and Figure 1 (B) shows application of SLM-Transformation on a string of integers. The output arrays in SLM-Transform are called *Ions Array (IA)* and *Buckets Array (BA)* which will be explained in detail in the following sections. Further, our technique constructs ion buckets with near uniform number of ions per bucket to further reduce the memory overhead for storage of sorted indexed ions. The ion buckets store only the information of piecewise ion frequency distribution in the bucket which can be used to approximately locate ions based on the mass of query fragment. Moreover, another technique employed by SLM-Index allows us to generate and store only a limited number of theoretical b-and y-ions from each peptide sequence which yield a higher probability of correct fragment-ion matches in open-search.

**Figure 1:**
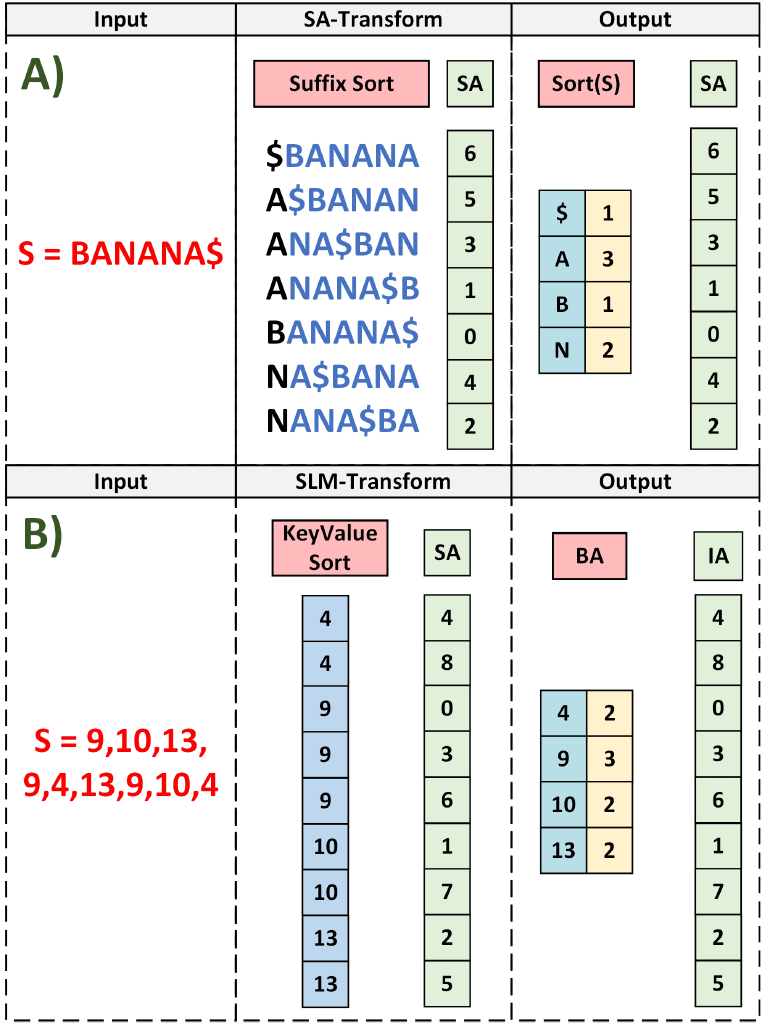
**A)** The input string S is transformed by Suffix Sort and the output is a compact version of sorted input along with Suffix Array that allows constant time querying of characters in S. **B)** The input integer string is transformed by Key-Value sort to allow a similar compact representation (BA) of input string as well as Ions Array (IA) which is analogous to SA.

The results show that our proposed method is highly memory efficient. We also show that our method allows constant-time queries. Moreover, the query speed is independent of index size and precursor mass tolerance window. SLM-Index provides open-source application programming interfaces (APIs) that can be easily utilized with most existing database search software.

## 2 The SLM-Transform Method

The prime transformation method employed by SLM-Transform sorts array indices using subsequent values as keys which is similar to suffix array construction for text data. This transformation is compounded by distribution of ions to buckets and further splitting the buckets in to smaller chunks using splitter ions. The distribution of ion frequencies between each splitter ion pair in a bucket is estimated using algebraic or statistical methods. The same estimation technique is then used to approximately calculate positions of indexed ions in a bucket when queried by mass.

Another major space reduction technique is based on the observation that if almost all peaks in its MS/MS spectrum have unknown mass shifts, then its correct sequence must have unknown modification at or near both termini. In this case, no fragment (peak) in that MS/MS spectrum would correctly match to the indexed ions even if a similar peptide sequence was present in the index. However, if the unknown mass shift locations in a MS/MS spectrum correspond to near center and/or at only one terminal of its correct peptide sequence, a fraction of peaks from N-, C-or both termini would match to correct b- and y-ions respectively in the index. This implies that in open-search where the search objective is to identify spectra with unknown mass shift, a large fraction of peaks in the spectra as well as indexed ions do not contribute to correct fragment-ion match scores. Based on this, SLM-Index limits the number of ions that are generated and indexed from N-or C-terminus of a theoretical peptide sequence. i.e. b-and y-ions respectively to F. The subsequent sections discuss SLM-Index construction in detail. The complete SLM-Transform based indexing and querying methods are shown in Figure 2.

**Figure 2:**
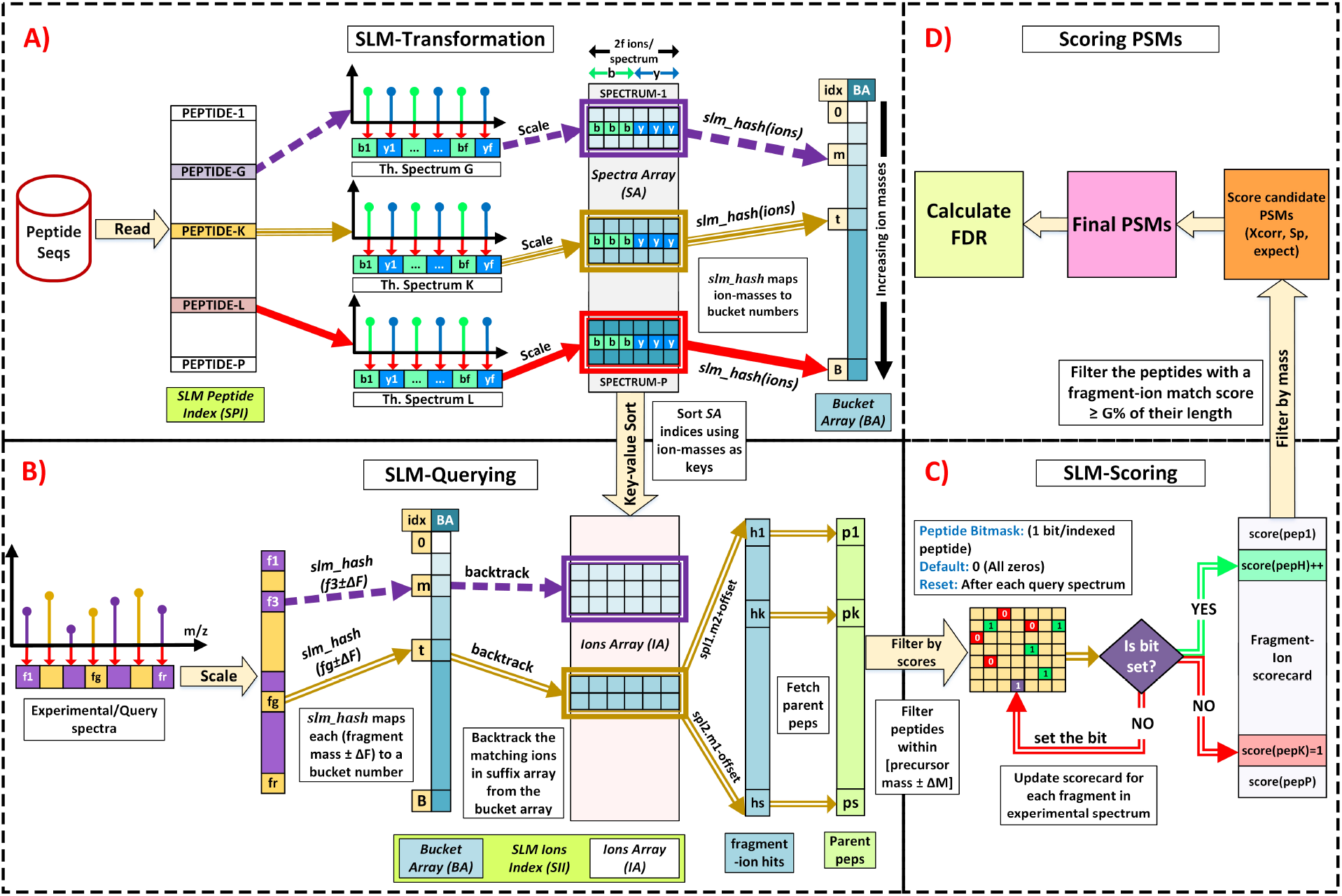
**A)** The peptide sequences are populated into SLM Peptide Index and corresponding theoretical spectra are generated and populated into Spectra Array. The SLM-Transform is applied on Spectra Array to produce Buckets Array (BA) and Ions Array(IA). **B)** The query fragments are first hashed into a bucket using *slm_hash.* The position of ions within the bucket that match the query ion are approximately calculated using hashed bucket information. **C)** The parent peptide IDs are calculated for fragment-ion matches (hits) and the scorecard entry for all hits are updated. The peptides with a fragment-ion match score ≥ G% of their length qualify as candidates which are further filtered based on precursor mass. **D)** The candidates are now scored which results in peptide-to-spectrum matches (PSMs). The PSMs can then be re-ranked and a confidence level is assigned using false discovery rate (FDR) algorithms.

### 2.1 SLM-Index Initialization

An empty SLM-Index is initialized by providing initial capacities for normal and modified peptides in index. SLM-Index capacities can also be automatically initialized by providing a peptide sequences file along with variable modifications information. A peptide sequence file can be generated from a proteome database using the methods and tools described in Section 7.

### 2.2 SLM Peptide Index construction

The normal (unmodified) peptide sequences along with their origin protein IDs, peptide-mass and sequence lengths are inserted into SLM-Index which are populated into an array called *pepArray* (*pA*). Once all normal peptide sequences have been populated, the *pepArray* is sorted by peptide-mass as shown in Figure 2. Assuming the total number of peptide sequences populated is denoted by *peps.* For each *pepArray* entry, the file-index is denoted by *fi* (4-bytes), peptide sequence by *seq* (64 bytes), length of peptide sequence by *len* (2 bytes), peptide mass by *mass* (4 bytes), and index of origin protein by *prot* (12bytes), the memory consumed by the *pepArray* can be analyzed as follows:

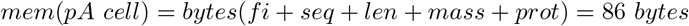

Hence,

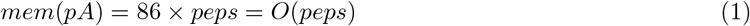

The modified variants of sequences in *pepArray* are inserted into SLM-Index by providing indices of their normal versions in *pepArray*, variant mass and modified sites information. This information is populated in an array called *modsArray* (mA). Once all variant peptides have been populated, the *modsArray* is sorted based on variant mass. Assuming the total number of modified peptides populated is denoted by *mods,* the normal peptide version index by *seqid* (4 bytes), variant mass by *vmass* (4 bytes), and *SLMvarAA* structure is denoted by vAA (12 bytes), the memory consumed by the *modsArray* can be analyzed as follows:

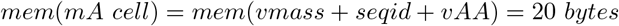

Therefore,

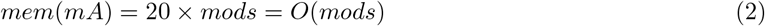

The *pepArray* together with *modArray* constitutes *SLM Peptide Index (SPI).* The SLM-Index can handle up to 10 modified residues per variant peptide and up to 7 types of modifications per index. The total memory consumption for *SPI* can be computed using (1) and (2) as follows:

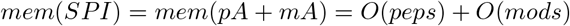

But we know,

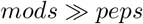

Therefore,

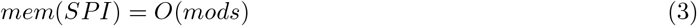

### 2.3 SLM Ion Index construction

For each peptide sequence in *SPI*, a theoretical spectrum is also inserted into the SLM-Index. The theoretical spectra at input are assumed to contain mass-to-charge (m/z) values of first *F* b- and first *F* y-ions up-to +3 charge state. Zeros must be appended in the generated theoretical spectra if the number of ions generated by a peptide sequence are less than 2F. Our empirical studies have shown that generating theoretical spectra with a F from 30 to 60 returns enough fragment-ion matches to filter potential peptide candidates. The b- and y-ions inserted for each theoretical spectrum are sorted by their m/z values and scaled to integers which are populated in an array called *Spectra Array (SA).* An empty array with B elements, called *Bucket Array (BA),* is created to store information on distribution of ion-mass frequencies in each bucket. The non-zero mass ions in Spectra Array are distributed among B buckets using a function called *slm_hash* to have near uniform distribution of ions within each bucket. Non-uniformly distributed ions might result in several false-positives in query results.

SLM-Index uses the bucket information to approximately locate bucketed ions when queried. For each bucket, we choose a set of splitter ions *(sp)* within the bucket and perform regression on the frequency of ions that lie between a pair of splitters. The location of a query ion in the bucket with respect to a splitter location is the sum of frequencies of ions that lie between splitter and query ion. This sum can be estimated by the area under regression polynomial in that interval. Note that the estimation can be also be done using a variety of sophisticated statistical and algebraic methods which are out of scope of this paper. An example of ion-frequency approximation using 1st degree polynomial and 1 splitter is shown in Figure 3. Each *Bucket Array* entry stores the location of first ion in bucket (loc), the maximum and minimum ion-masses in the bucket (*hi* and lo respectively), frequencies, masses and positions of splitter ions (*fsp, msp* and isp respectively), and the slope of regression polynomial on both sides of splitter ions (*m*1 and *m*2).

**Figure 3:**
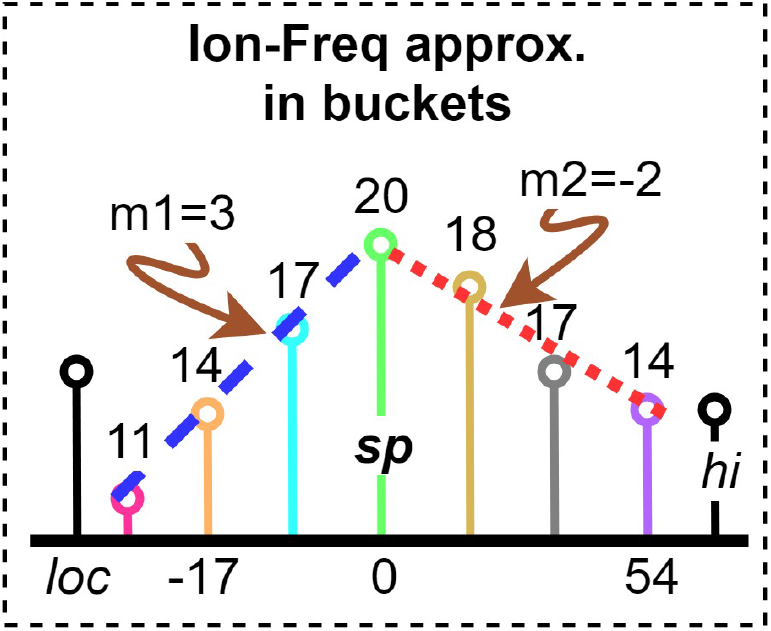
The ion frequencies in the sample bucket can be approximated using lines of slope (*m*) 3 and −2 towards left and right of splitter ion respectively. The query ion positions with respect to splitter ion position can be approximately calculated by area under the trapezoid formed between the two ions.

The *Index Array (IA)* is then constructed by applying SLM-Transformation. i.e. sorting the indices of *SA* using stored ion-masses as keys as shown in Figure 2 (A). After SLM-Transform, the *k^th^* entry in *IA* contains the index number of *k^th^* largest non-zero ion mass in SA. The *Index Array* along with *Buckets Array* constitute *SLM Ion Index (SII).* The *Spectra Array* is discarded from the memory after construction of *SII*. The upper-bound on *SII* memory is small as compared to other leading indexing strategies. This allows us to efficiently handle memory operations hence minimizing penalties that may arise due to several memory allocations/deallocations. Assuming the total number of peptides in SLM Peptide Index is denoted by *N*, the memory consumed by *SII* can be analyzed as:

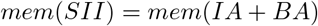

where:

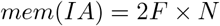

Assuming:

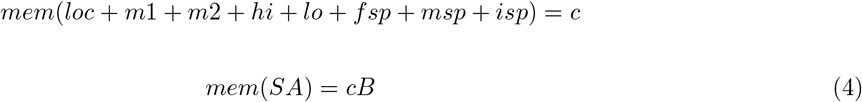

We have,

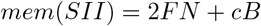

We know, *N* ≫ *B* and *F* is constant

Therefore,

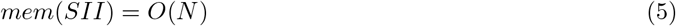

Figure 4 illustrates an example of SLM-Index construction.

**Figure 4:**
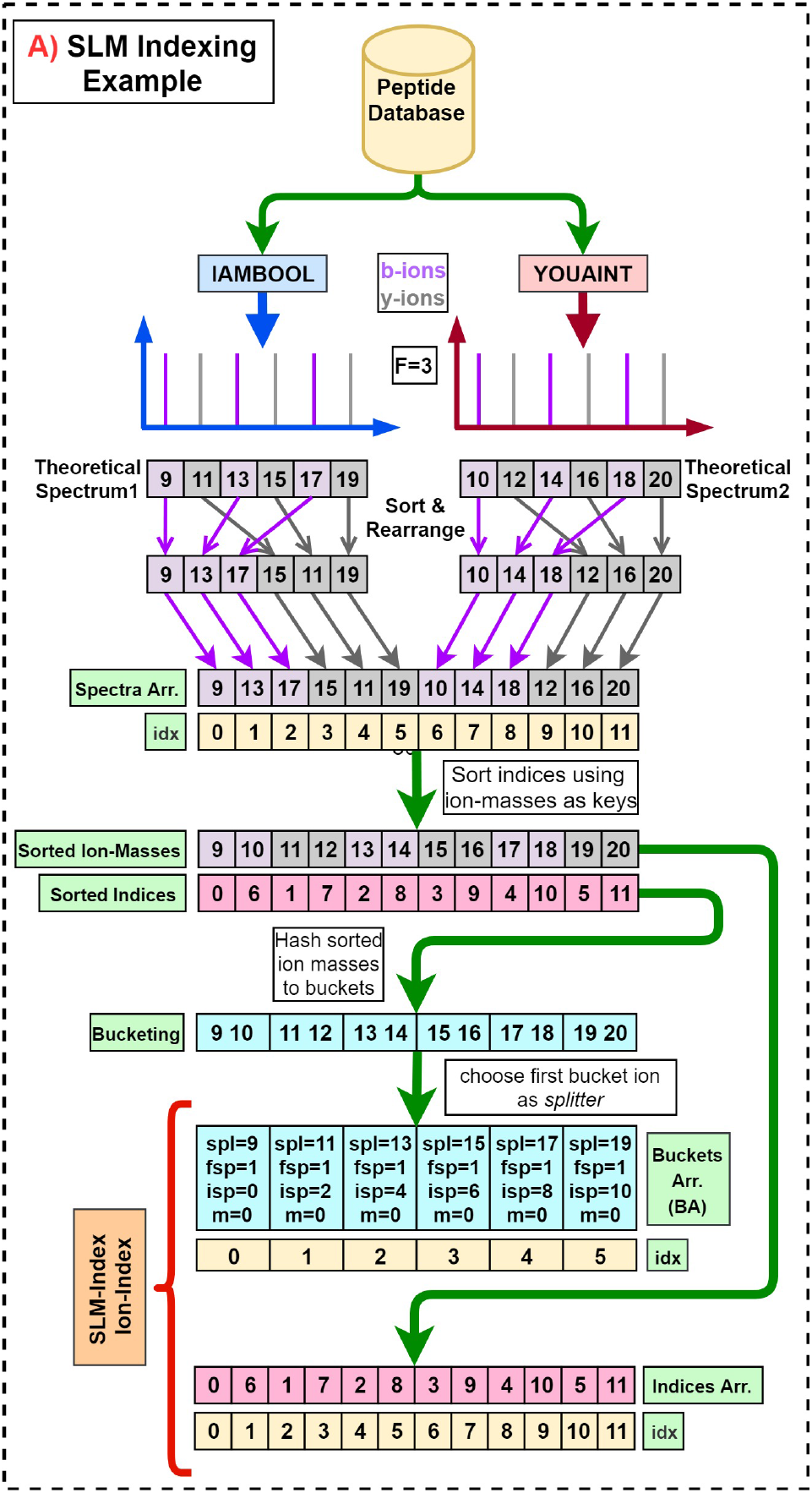
An example of SLM-Transformation using two peptide sequences. The b- and y-ions ions indexed for each peptide sequence are restricted to 3. The number of buckets is chosen to be 6 and the first ion in the bucket is chosen as the splitter ion. The slopes on left and right side of splitter are set to be 0 and 1 respectively.

### 2.4 Memory Analysis of SLM-Index

The total memory consumed by SLM-Index *(mem(SLM*)) can be calculated as:

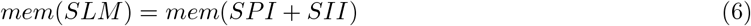

Using Equation (3) and Equation (5), we have,

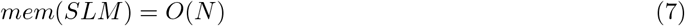

The memory consumption for SLM-Index peaks for a brief time to almost twice when *Index Array* is under construction and (equal size) *Spectra Array* has not been discarded from the memory yet. Therefore, the peak memory consumption for SLM-Index can be calculated as:

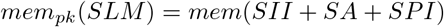

Using (6) and (4) we have:

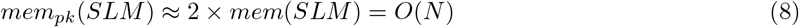

### 2.5 Time Analysis of SLM-Transform

SLM-Transform involves population of peptides (and variants) along with their spectra, hashing of ions to buckets and finally key-value sorting the *SA* indices. The time complexity for transformation is calculated by considering the time complexity of each construction stage. The peptide and variants can be populated in *O*(*N*), bucketing of ions can be done in *O*(*N*) as the total number of ions in the index are limited to *2FN*, and finally, the sorting can be done in *O*(*NlogN*). Therefore, the total indexing time (*Ti*) can be summarized as:

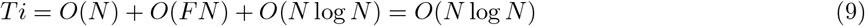

## 3 SLM-Index Querying

SLM-Index does not perform any preprocessing on input experimental MS/MS data. Experimental query spectrum is represented by an array of R (≥ 2*F*) elements, called *Query Array (QA)*, which contains the masses (m/z) of *R* peaks chosen from the input spectrum and inserted into *QA* by the user. Note that a wrong choice of peaks from the MS/MS spectrum may lead to poor PSM results. The SLM Query algorithm is explained as follows:

### 3.1 The SLM Query Algorithm

SLM-Transform based fragment-ion search begins by scaling each fragment mass-to-charge (m/z) in the *Query Array* to integer which is then hashed to a bucket number between [0-B] using *slm_hash* as shown in Figure 2 (B). The ions within that match the query fragment mass (± fragment mass tolerances) *(hits)* are approximately located by first determining a set of splitter ions (*sp*1 and *sp*2 respectively) in the bucket whose masses flank the query mass.

Then, the offset in position of query ion from each flanking splitter ion’s position is equal to the frequency of ions that lie in the mass interval between the splitter ion and query fragment (Δ). This summation is approximately equal to the area under the regression polynomial over this interval. Consider a query fragment having mass *frag*, the indexed ions that lie in the user defined fragment mass tolerance range, denoted by *hits,* can be located along with their parent peptide ids (*ppid*), ion series (*iser*), and ion series number (*inum*) in constant time using Algorithm 1.

For each fragment-ion match, the corresponding parent peptides scores are updated by *SLM Scorecard (SSC).* SLM-Index creates a 1-byte scorecard entry for each peptide in *SPI* and the *SSC* operations are governed by a bitmask contains one bit (clean/dirty) status for corresponding *SSC* entry. If the bitmask bit at *Parent Peptide ID (ppid)* is unset, that means that the value in scorecard is dirty and needs to be initialized for the first fragment-ion. In this case, the scorecard entry at the *ppid* is to intitialized to 1 and the bit is also set to mark the entry as clean. If the bit is already marked clean at *ppid*, then the score can be simply updated by +1 as depicted in Algorithm 2. The memory consumed by *SSC* can be analyzed as:

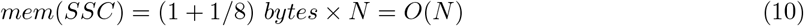

For each query spectrum, the search space is filtered in two stages, first based on fragment-ion match scores and second based on precursor mass tolerance window. The first stage filtration is done based on the fragment-ion scores (*Scr*) obtained by the peptides in the filtered range. Then the second stage filtration is done by first obtaining a peptide range by binary searching precursor mass of query spectrum against *SPI* and discarding the bitmask entries that lie outside of obtained range. A peptide, (*p*), is considered a potential candidate if its fragment-score is greater than or equal to the user-specified fraction (G%) of its sequence length (lsq). This is given by:

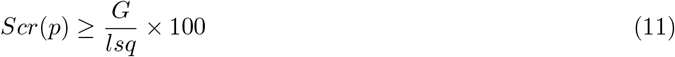

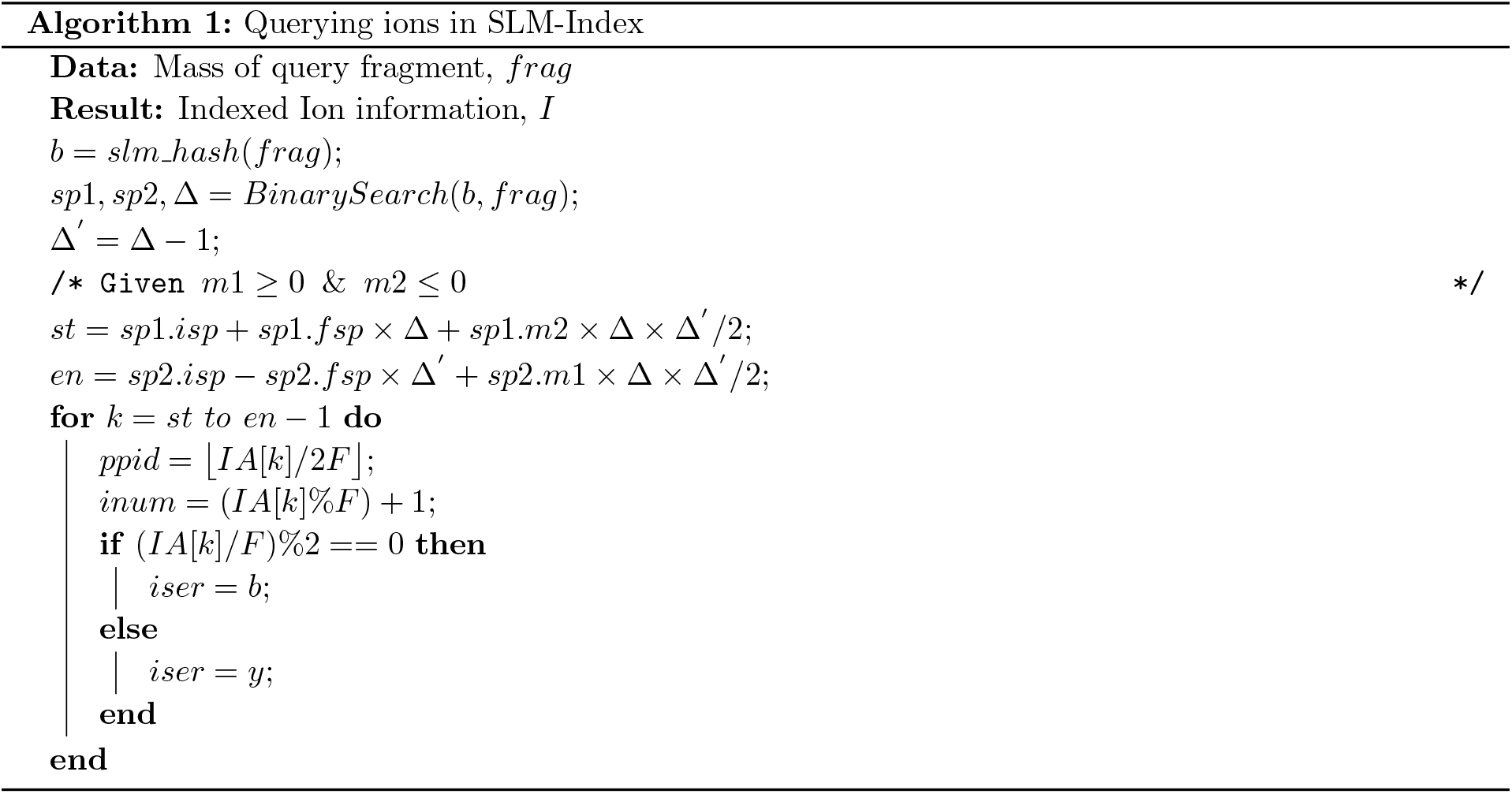

The only limitation with filtering candidate peptides using this technique is that it may fail to correctly spectra that correspond to peptide sequences having PTMs at or near both C- and N-termini. We studied the effects on search speed and sensitivity with varying *G* and our empirical studies showed that setting *G* between 9-13% (default: 10%) provides a good trade off between speed and sensitivity for most cases. The scoring algorithm is shown in Figure 2 (C and D). An example of querying fragments using SLM-Index is shown in Figure 5.

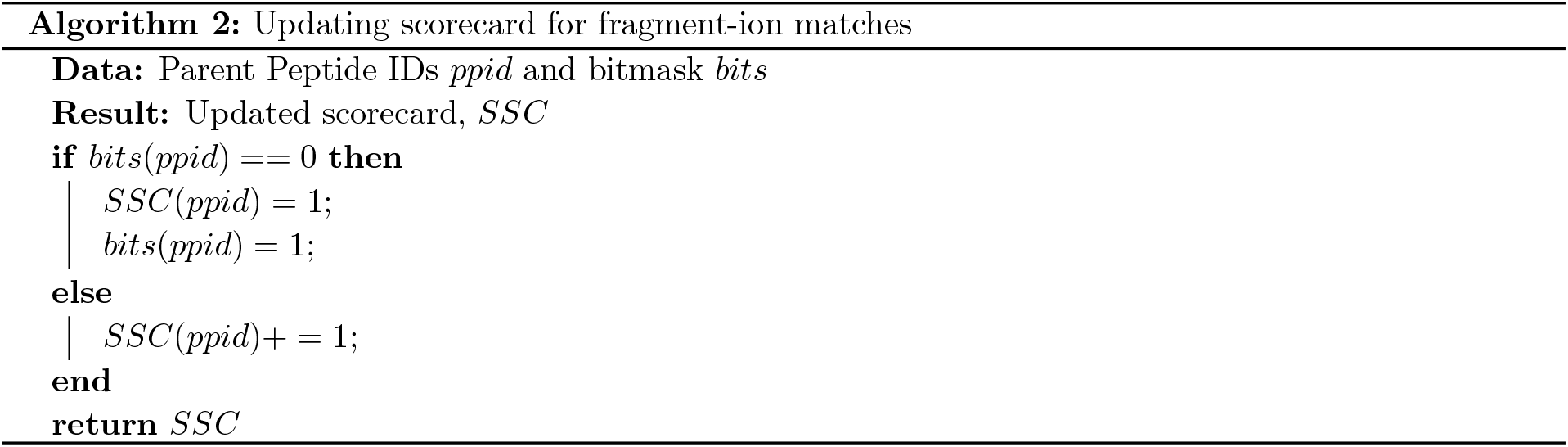

**Figure 5:**
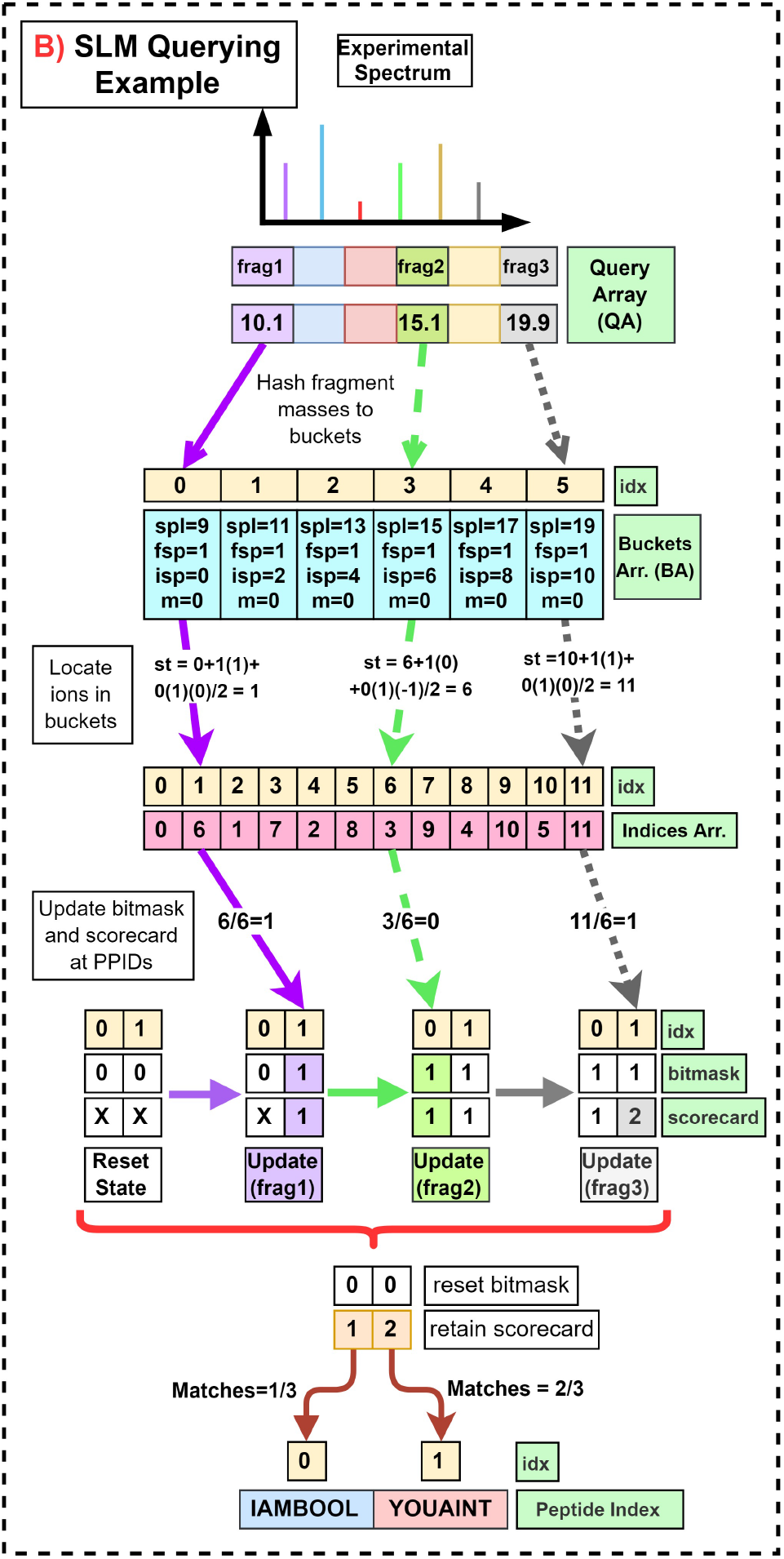
An example of querying 3 fragments from a MS/MS spectrum against the SLM-Index constructed in Figure 4. The query ions are hashed to BA and their locations in the bucket are calculated and the scorecard for parent peptides is updated. The PSMs can be deduced based on their fragment-ion scores.

### 3.2 Query Time Analysis

SLM-Index allows constant time query for each fragment-ion hit. However, the total search time for a SLM-Index fragment-ion database search depends on the average number of hits (*avh*) encountered per query fragment. *avh* is directly proportional to the precursor mass window, fragment mass tolerance and the number of peptides in the index and is inversely proportional to number of buckets and splitters chosen within each bucket. Assuming the number of hits per query fragment to be *avh,* the time for each query (*Tq*) and total search time for MS/MS spectrum (*Ts*) is computed as follows:

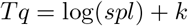

As the number of splitters *spl* is constant,

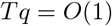

Therefore,

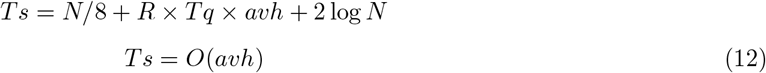

## 4 Results

The results discussed in the following sections have been obtained from the beta version of SLM-Transform which was integrated into the code base of Comet-MS v2016.03. The results are completely applicable for final version of SLM-Transform as the total size of the constructed index and query speed for both versions are the same. The beta implementation utilizes libdivsufsort (available at: https://github.com/y-256/libdivsufsort) [22] for key-value sorting of ions to construct the *IA.* The beta version has been tested with Comet-MS Ion Trap MS/MS resolution setting which scales the ions by 1/1.0005 ≈ 0.99*Da*. The low-resolution scaling factor settings limits minimum fragment-ion tolerance to 1Da for ion-bucketing. The beta SLM-Transform implementation is available at: https://github.com/pcdslab/slmindexbeta

### 4.1 Memory Footprint Comparison

We compared the memory consumption for SLM-Index against MSFragger by creating indices of various sizes using using publicly available proteome databases. The index size comparison for SLM-Index was made against MSFragger GUI v6.0 running MSFragger-20180316 and Philosopher 20180317. The databases employed for our experiments were Homo Sapiens (UP000005640) and Rattus Norvegicus (UP000002494) appended with reverse decoys and cRAP MaxQuant. The indices of increasing sizes were generated by varying the number and types of modifications added to the index. The following digestion parameters fixed for all experiments: Digestion=Tryptic; Allowed Peptide Lengths= 3 to 50; Allowed Missed Cleavages: 2; Digested Mass Range: 500 to 5000 amu; Duplicate Peptide Sequences=No.

The memory consumption trend for both SLM-Index and MSFragger against increasing number of indexed peptides is shown in Figure 6. The index size was increased till the point when SLM-Index completely exhausted the available system memory as the current implementation version does not yet support splitting and caching its index onto the disk.

**Figure 6:**
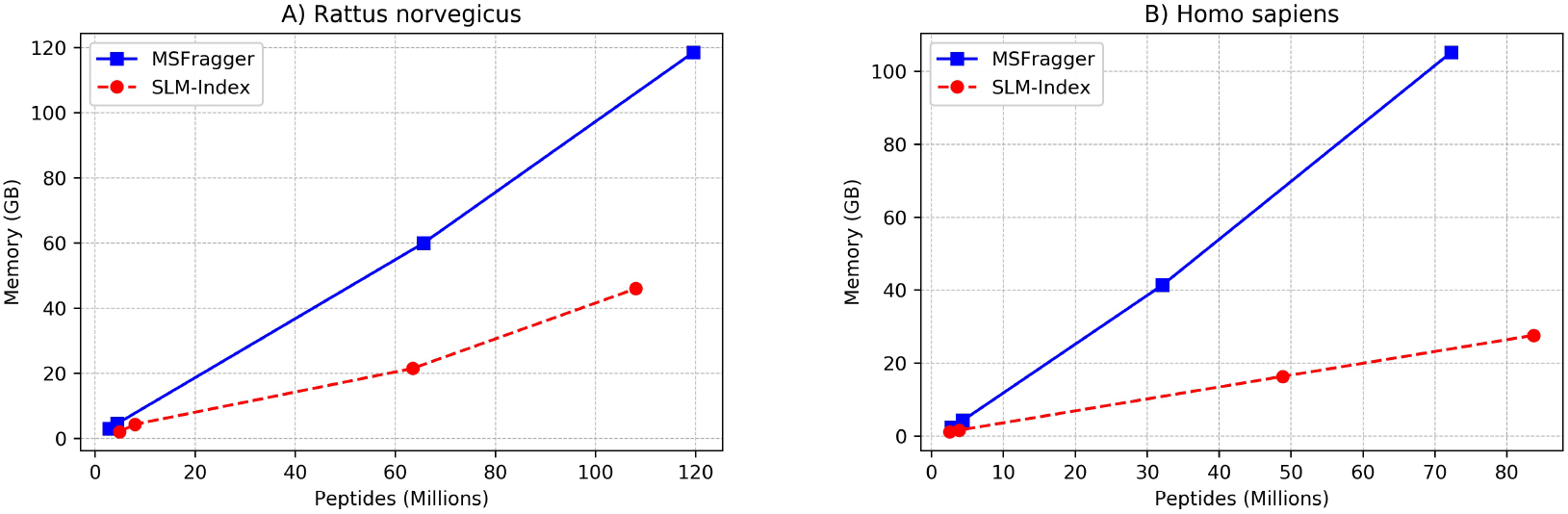
The memory footprint comparison between SLM-Index and MSFragger index for **A)** Rattus norvegicus and **B)** Homo sapiens database with increasing number of peptides in the index (generated varying number of modifications in the index) shows significantly small and highly linear growth rate for SLM-Index.

### 4.2 SLM-Index allows constant time queries

The query time for SLM-Index was evaluated for varying index sizes and number of queries performed on the index. The experiments were conducted on a server machine with 24 cores of Intel®; Xeon®; CPU E5-2620 v3 @ 2.40GHz, having 48GB RAM, running Ubuntu 14.04.02. The SLM-Index size was varied from 27 to 92 million peptides and the experiment was repeated several times for each index size. Figure 7 shows the SLM-Index query speed with increasing index size.

**Figure 7:**
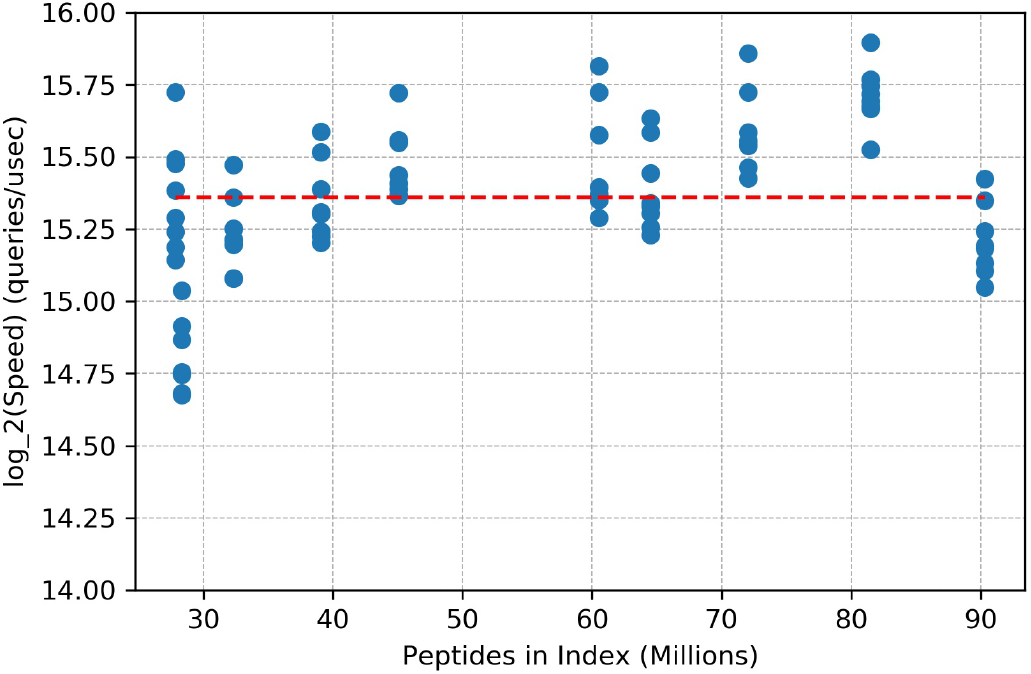
The query time trend is near constant (shown by dotted red line) with increasing index size. Note that the precursor mass tolerance window size was varied for each experiment but no significant change was observed.

The query speed for SLM-Index was evaluated by searching 60 experimental spectra from UPS2 dataset against a rat database combined with several combinations of modifications for variable index sizes. The search space was expanded by setting the fragment mass tolerance to 1*Da* and precursor mass-tolerance window to 100Da. The results indicated a near constant query speed of 41 *queries/ns.* Note that the query speed is independent of index size or the precursor mass tolerance window size. However, the total time to search a query MS/MS spectrum depends on the number of query hits encountered by SLM-Index. Please refer to *Supplementary Table 5* for detailed query speed results.

## 5 Discussion

We studied the accuracy of the PSMs reported by SLM-Transform based Comet-MS under different settings. The search accuracy was calculated as the fraction of correctly identified PSMs when compared against original ground-truth sequences. The search accuracy trend against each parameter under study was obtained keeping the rest of parameters and conditions constant. We further combined all the acquired trends to study the combined effect of all the parameters together. The details on datasets along with the digestion settings for the database used for following experiments can be found in *Methods* section.

### 5.1 Varying Number of Modifications added to Index

We studied the impact of varying the number of commonly known modifications added to SLM-Index in this section. In the first set, CID/HCD dataset was searched against Rattus norvegicus database varying (0 to 3) number of modifications added to the index keeping precursor mass window fixed. The same experiment set was then repeated with varying settings of precursor mass window as well. The results showed that over more than 90% of spectra in CID/HCD unmodified dataset were correctly identified. However, the identification rate for modified CID/HCD and simulated datasets increased significantly as as we added upto 2 modifications in the index and started saturating after that. To validate the above observation, the simulated dataset was searched against the index with number of modifications varying from 0 to 3. The results indicated an expected increasing trend in the identification rate as the number of modifications added to the index were increased. The experiment was repeated for different window widths as well. The above experiments imply that the percentage correct identification rate for a MS/MS dataset increases as the number of similar or correct peptide sequences are increased in the index. Based on the obtained results, we conclude that carefully adding a few but abundantly present modifications at fairly wide window sizes yields optimum identification rate in contrast to blind or narrow-window search with all 2*^Mods^* combinations. Figures 8 (A, B, C) show the trend in identification rate with increasing number of modifications added across different window settings. Also, the trend of identification rate with respect number of modifications added for different window sizes is shown in Figure 10 (A, B, C).

**Figure 8:**
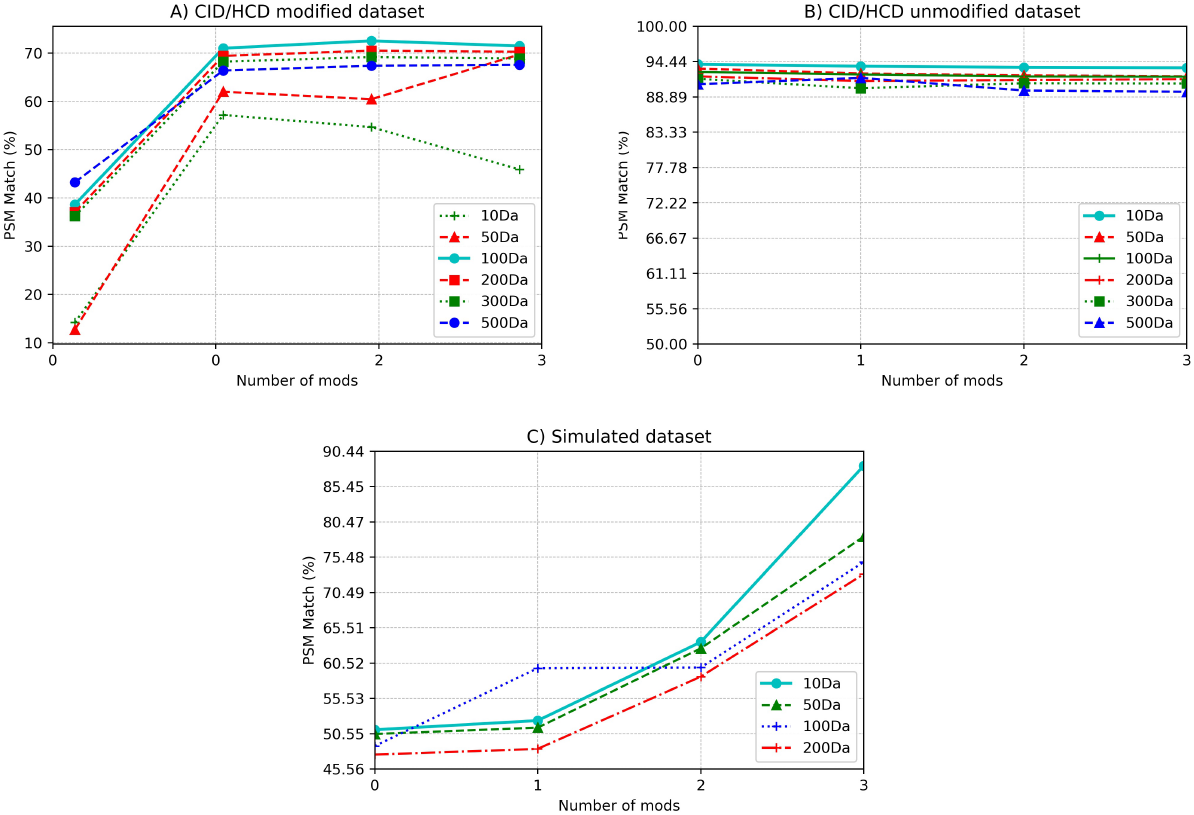
**A)** The identification rate for the CID/HCD modified dataset increases as the number of modifications added to the index are increased from 0 to 2 and then saturates for all the window sizes except 10 Da. All unknown modifications have masses > 10Da hence it fails to identify any unknown modifications. **B)** The identification rate for the CID/HCD unmodified dataset remains almost constant as the normal peptides are included in the index in all cases. Please note that the data point for 3 mods and 500 Da shown in this plot was extrapolated and not acquired experimentally. C) The identification rate for the simulated dataset increases significant when the number of modifications added to the index are increased from 0 to all 3 modifications in the simulated data set.

### 5.2 Varying Precursor Mass Tolerance Window Width

We studied impact of variation in the precursor mass tolerance search windows in this section. The experiments were performed on both CID/HCD and simulated datasets. The search windows were varied from 10Da to 500Da in all sets of experiments. We repeated this set of experiments with different number of modifications added to the index. The search results for unmodified CID/HCD dataset were almost constant in all experiments with a small 4% variation in results over wide window sizes. However, the fraction of correctly identified PSMs for modified CID/HCD and simulated datasets improved significantly as the window size was increased from 10Da to 100Da and then saturated.

We observed that the correct identification rate is optimum if the search window width is set in the range of sum of unknown modification masses in the dataset. Expanding search window sizes beyond unknown mass ranges incurs a deterioration in accuracy as the probability of hitting a false-positive increases significantly. To test this hypothesis, we performed similar experiments using the simulated dataset. We noticed that the optimum results were obtained at narrow-windows as the sum of unknown modifications in dataset was less than 100Da. Figures 9(A, B, C) show the trends in the identification rate against varying window widths with various number of mods settings.

**Figure 9:**
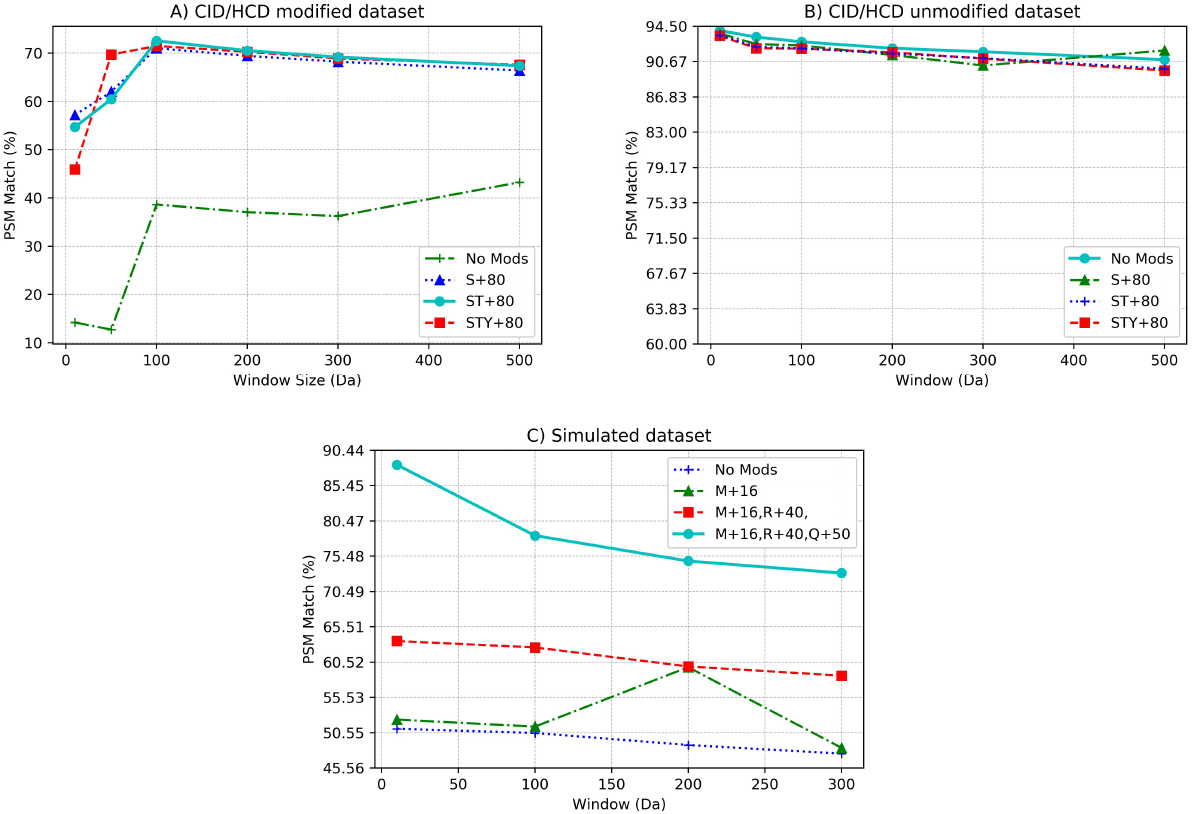
**A)** The optimum identification rate for the CID/HCD modified dataset was obtained between window size of 100 to 200Da when at least one modification is added to the index and between 300 to 500Da in case of blind search indicating presence of unknown modifications in corresponding windows. **B)** The identification rate for the CID/HCD unmodified dataset remains almost almost the same. Please note that the data point for 3 mods and 500 Da in this plot was extrapolated and not acquired experimentally. **C)** The optimum identification rate for the simulated dataset is achieved in window ranges between 10 Da to 100Da and when more modifications are added to index.

### 5.3 Varying the peptide coverage of simulated data

In the following set of experiments, we studied the effects of the input MS/MS data quality on the rate of correctly identified PSMs. The entire CID and HCD dataset was simulated using the available ground-truth sequences by using MaSS-Simulator. The maximum charge was set to 3, signal-to-noise ratio was set to 2.8 mass-range was set to 100Da to 7000Da and the rest of the parameters were kept default apart from the peptide coverage, which was varied from 40% to 100% with an increment factor of 10%. We searched the 7 generated datasets against RAT database in optimum settings i.e. precursor mass tolerance window set to 120Da and phosphorylation on serine (S) and threonine (T) residues added to the index. The trend in correct identification rate against variation in peptide coverage can be seen in Figure 11. The search results were predictable and the correct PSM match percentage increased as the peptide coverage was increased. The increasing trend of percentage correct PSM match against the peptide coverage is shown in Figure 11. See the *Supplementary Table 4* for detailed results.

**Figure 10:**
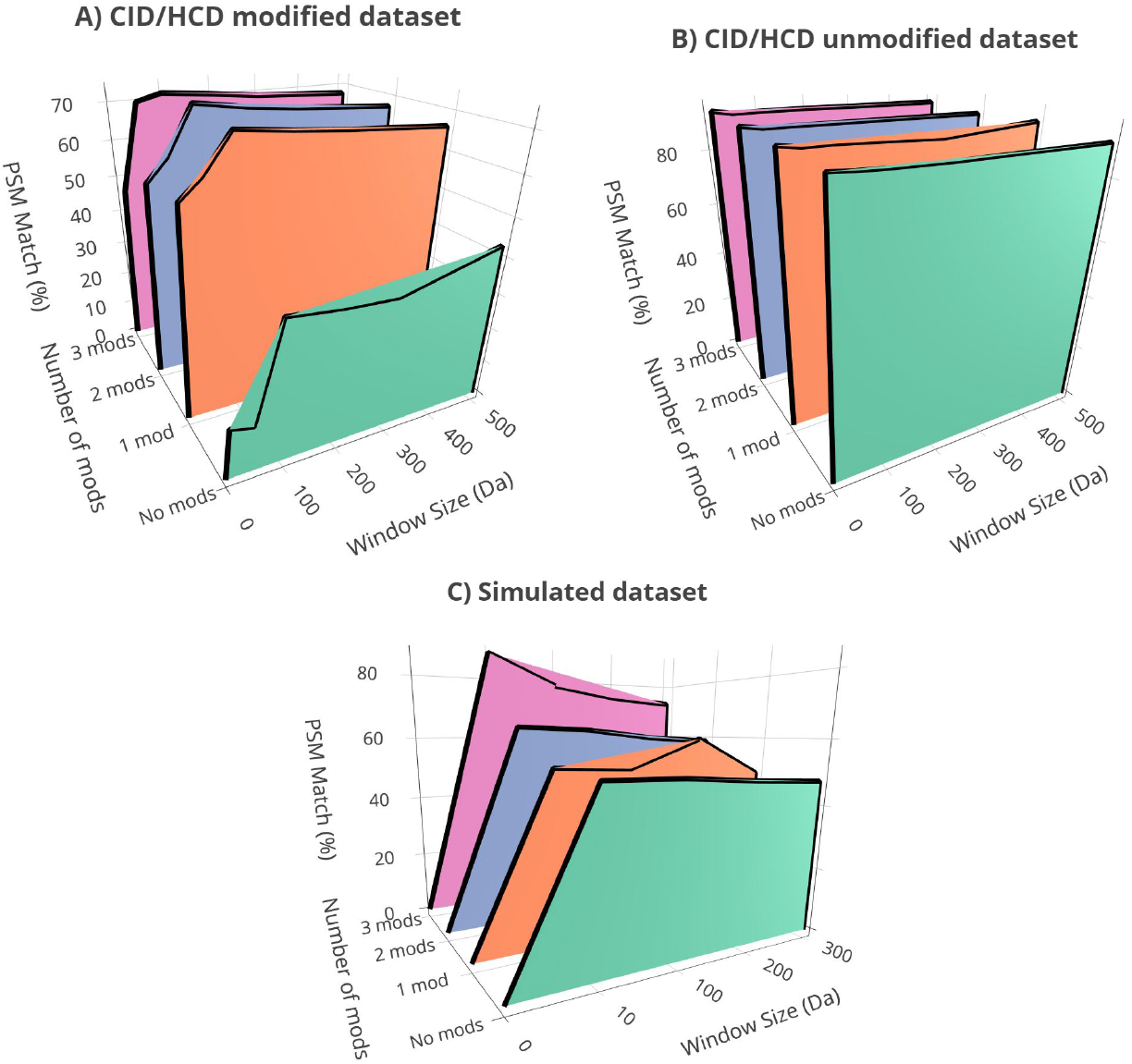
**A)** The results indicate an abundant presence of phosphorylation at S and T residues along with other unknown modifications within 150Da in the CID/HCD modified dataset. **B)** The results indicate a high identification rate for any number of modifications added or window size indicating an abundance of unmodified peptides in the CID/HCD unmodified dataset. Please note that the data point for 3 mods and 500Da in this plot was extrapolated and not acquired experimentally. **C)** The results indicate presence of unknown modifications within 100Da window in the simulated dataset.

**Figure 11:**
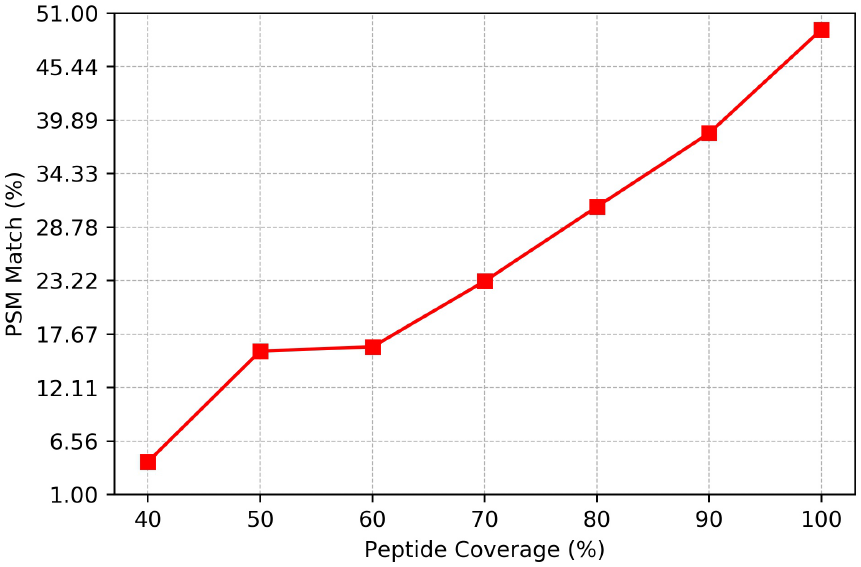
The identification rate for the simulated dataset increases significantly as the quality of the simulated data set is improved. The results show that increasing peptide coverage yields substantial improvement in correct identification rate.

### 5.4 Summary of results

The combined results are inline with our initial hypothesis that running a blind search over extensively wide precursor mass tolerance windows results in a significant loss in sensitivity, becomes prone to false positives and incurs search time penalties. On the other hand, searching over narrow windows taking a huge variety of known modifications in the index results in an exponential expansion in index size along with the fact that the spectra with unknown modifications still escape the search space. This implies that an optimum search accuracy is obtained when the search is run in two phases. i.e. first phase with the maximum modifications added to the index in a narrow-window mode, second phase with only the expected abundant modifications added to the index in a mildly open-window. See the *Supplementary Tables 1, 2 and 3* for detailed corresponding results. To summarize the findings:

1. High quality MS/MS datasets yield better correctly identified peptide matches.
2. The correct identification rate improves as the number of modifications considered in the index are increased.
3. The precursor mass window widths should not be widened too much beyond expected range of sum of mass of unknown modifications in the dataset.

## 6 Implementation

SLM-Transform has been implemented in C++ as an API based library to allow abstractions which can be used to integrate SLM-Index within any other tool. Moreover, we implemented error checks at multiple locations in our code base to gracefully return errors and avoid a fatal crash. SLM-Transform can be built using Linux®; GCC toolchain or MinGW based GCC toolchain on Windows®; machine. The Wiki available on SLM-Transform GitHub details all the required tools and steps for building and using SLM-Transform and SLM-Transform beta which has been integrated into Comet-MS codebase. Some of the limitations in our current implementations are summarized as follows:

1. The current implementation of Buckets Array only supports 1 splitter ion per bucket viz. the ion having maximum ion frequency in the bucket.
2. The regression polynomial of degree 1 is used for ion location approximation in a bucket. The approximation function is however exposed to public interface, so the users can configure according to their data. Moreover, a number of statistical and algebraic methods will also be made available within the library in future releases.
3. The current implementation constructs *BA* using *direct-hashing* based *slm__hash* function.

## 7 Methods

In the following sections, we describe the tools and methods used to preprocess proteome databases, and simulate and convert MS/MS data for our experiments.

### 7.1 Digested Peptide Sequences

Publicly available proteome databases from UniProtKB were used in our experiments. The proteome databases were digested in-silico into corresponding peptide sequence databases using Digestor.exe tool available with OpenMS v2.1.0 [30] or Protein Digestion Simulator [26]. Then, the redundant peptides sequences were removed using DBToolkit v4.2.4 [19] invoked with default parameters resulting in a non-redundant peptide database file. We use this file for any experiments performed and described in this paper.

### 7.2 Mass-Spectrometry (MS/MS) data simulation

The simulated MS/MS datasets used in our experiments were generated using MaSS-Simulator [1]. Several simulation parameters and post-translational modification settings were varied across experiments to generate a variety of ground-truth data. More details about each simulated dataset are provided with each experiment’s description.

### 7.3 MS/MS data format conversion

The MS/MS data, if in a different format, was pre-converted to mzML format before using in any of our experiments using msconvert.exe [14] tool available with ProteoWizard.

### 7.4 Dataset and Database Details

The following sections provide details on the datasets that were used for discussion experiments along with the settings used to digest the Rat database.

#### 7.4.1 CID and HCD datasets

The ground-truth CID and HCD datasets were obtained from [27]. Both the CID and HCD datasets have been experimentally (LC-MS/MS pipeline) generated from a rat sample. The datasets contain two subsets of ground-truth spectra. i.e. modified dataset which contains spectra that contain variable oxidation on methionine residues, phosphorylation on serine, threonine and tyrosine residues, and deamidation on arginine and glutamine residues. The unmodified dataset contains spectra that have no modification.

#### 7.4.2 Simulated Dataset

The simulated dataset was generated using Mass-Simulator and contains randomly chosen 1000 peptide sequences from Rattus norvegicus peptide database. The extracted sequences were modified with 3 custom modifications and the peptide coverage was set to 80% for simulation. The simulated dataset was then searched against the index with number of modifications varying from 0 to 3.

#### 7.4.3 Database Digestion Settings

The Rattus norvegicus database was downloaded from UniProtKB and was tryptically digested with 2 allowed missed cleavages, peptide lengths varying from 7 to 50 and peptide masses ranging from 100 to 7000Da.

## Acknowledgments

This research was supported by National Institute of General Medical Sciences (NIGMS) of the National Institutes of Health (NIH), United States under Award Number R15GM120820, and National Science Foundations (NSF) under Award Numbers NSF CRII CCF-1464268, NSF CRII CCF-1855441, and NSF CAREER ACI-1651724. The content is solely the responsibility of the authors and does not necessarily represent the official views of the National Institutes of Health or that of National Science Foundation.

## Conclusion

In this paper, we presented a scalable and memory efficient indexing method called SLM-Transform for fragment-ion based database search. SLM-Transform bounds its index memory by using both the elements and indices of its data structure arrays or information storage and retrieval. Moreover, the upper bound on memory allows for efficient memory allocations to avoid any performance degradation due to memory fragmentation. The results showed upto 4x memory footprint improvements for SLM-Index over MSFragger index. Further, fragment-ion based filtering may be complemented with other filtration methods such as sequence-tagging for even faster and refined search results.

Identification of peptides in a proteolyzed complex protein mixture by searching MS/MS spectra obtained from a mass-spectrometer is a critical step in proteomics studies. Experimental MS/MS spectra often have unexpected mass shifts due to PTMs and require an open-window in order to be captured by the search algorithm. Running a blind search with wide search window results in sensitivity loss. We show this by studying the impact of varying precursor window and the number of modifications added to the index on search accuracy. To improve the search accuracy, peptide variants with commonly occurring PTMs should be accommodated in the index which results in exponential increase in the index memory size which may hamper the search algorithm performance. Finally, conventional target-decoy based FDR filtering techniques are not recommended for open-search and hence, filtration techniques that consider unknown modifications while re-ranking must be employed.

